# Loss of ClpP Function in *Clostridioides difficile* 630 Significantly Impacts Sporulation Systems

**DOI:** 10.1101/2021.02.05.429978

**Authors:** Catherine E. Bishop, Tyler Shadid, Nathan P. Lavey, Megan L. Kempher, Nagib Ahsan, Jimmy D. Ballard, Adam S. Duerfeldt

## Abstract

The Gram-positive bacterium *Clostridioides difficile* is a primary cause of hospital-acquired diarrhea, threatening both immunocompromised and healthy individuals. An important aspect of elucidating mechanisms that drive *C. difficile* persistence and virulence relies on developing a more complete understanding of sporulation. *C. difficile* sporulation is the single determinant of transmission and complicates treatment and prevention due to the chemical and physical resilience of spores. Hence, the identification of potentially druggable targets that significantly attenuate sporulation is important. In this report, we describe the impact of the loss of caseinolytic protease P (ClpP) isoforms in *C. difficile* strain 630 on sporulation phenotypes. Using CRISPR-Cas9 nickase mediated genome editing, stop codons were inserted early in the coding sequence for *clpP1* and *clpP2* to generate *C. difficile* mutants that no longer produced ClpP1 or ClpP2. The data show that these genetic modifications lead to altered sporulation phenotypes, germination efficiencies, and cytotoxicity. Comparative proteome profiling of *C. difficile* 630 WT and *clpP* mutants reveals potential proteolytic targets of ClpP that are involved in sporulation. These analyses further reveal the potential for preferred co-chaperone interactions for each ClpP isoform. Taken together, our results demonstrate that ClpP, a promising target in other Gram-positive pathogens, holds promise as an anti-sporulation target in *C. difficile*.

## Introduction

*Clostridioides difficile* is a Gram-positive, anaerobic, spore forming bacterium and is classified by the Centers of Disease Control as an “urgent” threat to human health (Lawson, Citron et al. 2016, Prevention 2019). *C. difficile* infection (CDI) accounted for 223,900 U.S. hospitalizations in 2017, and was responsible for >36% of the total deaths resulting from antibiotic-resistant infections (Prevention 2019). A complicating factor in clinical treatment is that traditional therapeutics for CDI also eliminate host microflora, promoting gut microbiota imbalances, and making the host susceptible to further infection and/or CDI recurrence (Ananthakrishnan 2011). Another major obstacle to CDI eradication is the ability of *C. difficile* to form spores that are resistant to host clearance, routine physical sterilization techniques, and antibiotics (Dyer, Hutt et al. 2019, Mendo-Lopez, Villafuerte-Galvez et al. 2020). CDI transmission is dependent on the production of these aerotolerant spores, thus making anti-sporulation approaches a promising strategy worth exploring to aid in the mitigation of CDI (Lawley, Clare et al. 2009, Crobach, Vernon et al. 2018).

Until recently, the molecular drivers behind spore formation and germination remained relatively uncharacterized. Because aspects of sporulation were thought to be generally conserved among endospore-forming bacteria, information gleaned from *Bacillus subtilis* has historically served as a surrogate for studying sporulation in *C. difficile*, an organism that requires more stringent methodologies to handle and manipulate. The last decade, however, has capitalized on new methodologies and technologies allowing for the study of sporulation biology in *C. difficile* itself, revealing the critical transcriptional regulators of sporulation and providing insight into factors contributing to spore morphology (Shen 2020). While *C. difficile* maintains similarities to *B. subtilis* sporulation machinery, these new discoveries have helped highlight some discrepancies, as well. Likewise, the correlation between *B. subtilis* and *C. difficile* biology provide a compelling reason to interrogate systems that lie in the axis of sporulation of one organism in the other to continue to deconvolute the similarities and differences in sporulation biology.

As a key regulator of pathogenicity in infectious bacteria and given its roles in mediating protein turnover and bacterial homeostasis, caseinolytic protease P (ClpP) has emerged as a new target for antimicrobial development. The role of ClpP in *B. subtilis* was initially described by Msadek and Dartois over two decades ago (Msadek, Dartois et al. 1998). Subsequent studies have shown that the ClpP system regulates several pathways that influence sporulation and virulence in *B. subtilis* with distinct sporulation initiation pathways being controlled by ClpP and its cochaperones, ClpX and ClpC (Nanamiya, Ohashi et al. 1998, Liu, Cosby et al. 1999, Nanamiya, Takahashi et al. 2000). For instance, complexation of ClpP and ClpX (ClpXP) stimulates early-stage sporulation through degradation of the checkpoint polypeptide, Sda, which inhibits sporulation in response to DNA damage (Ruvolo, Mach et al. 2006). Complexation of ClpP and ClpC (ClpCP), however, initiates early forespore maturation through the degradation of SpoIIAB and subsequently indirect activation of SigF (Pan, Garsin et al. 2001).

While *C. difficile* harbors proteolytic machinery similar to that of *B. subtilis*, various orthologs (e.g., Sda) are missing from the *C. difficile* genome. Additionally, while the RNA polymerase sigma factors and sporulation regulatory proteins (Spo proteins) are present, significant mechanistic divergence of these factors from *B. subtilis* has been identified (Fimlaid, Bond et al. 2013). Thus, although the general roles of ClpP in bacteria seem to be conserved, the regulatory and signaling networks that depend upon ClpP are organism-dependent and have not been defined in *C. difficile*. Evidence that *C. difficile* sporulation biology diverges significantly from *B. subtilis*, provides a convincing justification for more detailed exploration of the ClpP system and involvement in sporulation biology in *C. difficile* itself (Fimlaid, Bond et al. 2013, Edwards and McBride 2017, Zhu, Sorg et al. 2018).

Contrary to *B. subtilis*, which contains a single isoform of ClpP, genes encoding two isoforms, *clpP1* and *clpP2*, are present in the *C. difficile* genome. These genes are expressed in exponential and stationary *C. difficile* growth phases, suggesting that both maintain active roles over the bacterial life cycle, though *clpP2* transcripts are observed at significantly lower levels compared to *clpP1* (Lavey, Shadid et al. 2019). This difference in expression and capability to operate as uncoupled proteases suggests a unique mechanism in which ClpP1 is the major contributor to cellular homeostasis, while ClpP2 may function in a more accessory role outside the bounds of general homeostasis (Lavey, Shadid et al. 2019). Additionally, proteomic analysis identified ClpP1 as part of the mature spore proteome (Lawley, Clare et al. 2009) and has linked ClpP1 and the cochaperones ClpC and ClpX with stress response in *C. difficile* 630 (Jain, Graham et al. 2011). The amino acid sequences of both ClpP1 and ClpP2 are highly similar to *B. subtilis* ClpP (74% and 63% amino acid sequence similarity, respectively) and there are no obvious differences between the known regulatory domains (Lavey, Shadid et al. 2019). Given the aggregate parallels yet divergent nature of sporulation mechanisms in *B. subtilis* and *C. difficile*, we hypothesized that while the ClpP system in *C. difficile* would be a key contributor to sporulation, new information about sporulation is likely to be learned by interrogating the effects of ClpP disruption in *C. difficile*. To begin our investigation, we report the essentiality of ClpP1 and ClpP2 on *C. difficile* growth, toxin production, and sporulation. We further demonstrated the impact of ClpP1 and ClpP2 absence on the *C. difficile* global proteome expression profile.

## Results and Discussion

### Construction of *clpP* mutants in *C. difficile* strain 630 (WT)

To determine if *clpP1* (CD630_3305) and/or *clpP2* (CD630_3351) is important for *C. difficile* growth, toxicity, and sporulation, we inserted stop codons 13 and 11 amino acids, respectively, downstream of the translation start site of the genes using a CRISPR-Cas9 nickase system (detailed in Materials and Methods and shown in **Figure 1**). PCR amplification of either *clpP1* or *clpP2* combined with *Nhe*I restriction enzyme digestion of PCR products revealed the presence of the inserted stop codon (**Figure 1B**). Sanger sequencing confirmed the presence of the desired stop codon in *clpP1* and *clpP2* (**Figure 1C** and **1D**, respectively) and whole genome sequencing confirmed that no off-target mutations were generated. Whole-genome sequencing data are available in the NCBI SRA database under BioProject identifier (ID) PRJNA675083.

**Figure 1.**
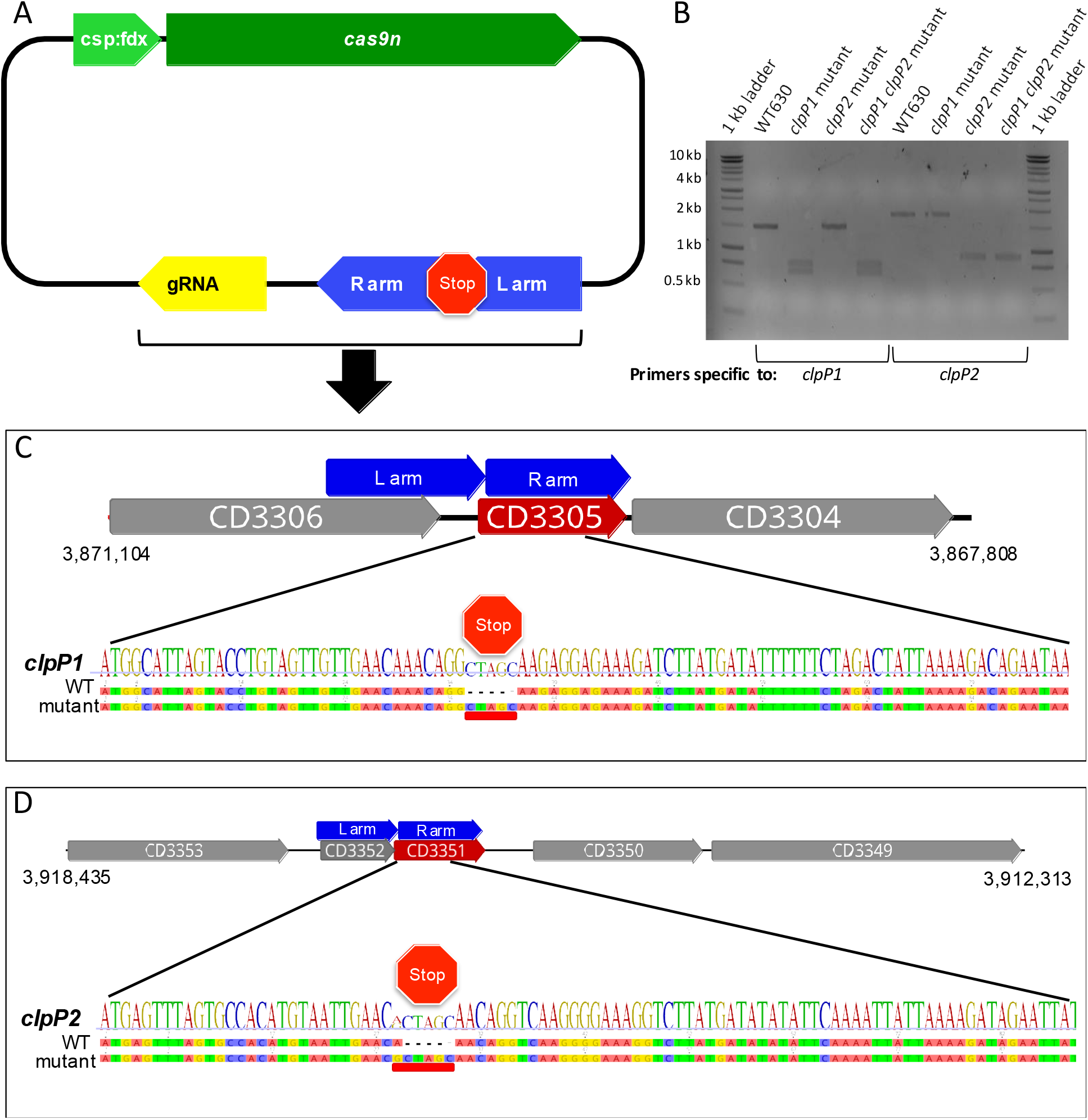
Genomic context of *C. difficile* strain 630 *clpP1 and clpP2*. (A) Cas9n vector map for the insertion of a stop codon in *clpP1* and *clpP2* generated on the pMTL84151 backbone. The modification of the backbone includes codon optimized *cas9* nickase genes, gRNA, and homology regions (left and right arms). (B) PCR amplification and NheI digest of *clpP* mutants compared to WT. (C-D) Gene maps of *clpP1* (C, top) neighboring genes (CD630_3304, *clpX*; CD630_3305, *ClpP1*; CD630_3306, *tig*) and *clpP2* (D, top) neighboring genes (CD630_33490, exosporium glycoprotein *bclA3*; CD630_33500, putative glycosyl transferase family 2; CD630_33520, AraC family transcription regulator; CD630_33530, GntR family transcriptional regulator) with homology arms indicated in blue. Sanger sequencing showed the insertion of a stop codon compared to unedited sequences 35 bp downstream of *clpP1* start site (C, bottom) and 28 bp downstream of *clpP2* start site (D, bottom).

### ClpP isoforms play a consequential role in *C. difficile* growth and toxicity

*B. subtilis* displays growth-deficient phenotypes when lacking ClpP function (Gerth, Krüger et al. 1998), which led us to examine the impact of ClpP1 and ClpP2 loss on *C. difficile* growth under typical and heat stressed conditions. Similar growth profiles were observed between WT and all mutant *C. difficile* 630 strains at 37°C (**Figure 2A**) and 42°C (**Figure 2B**). At both temperatures, however, the maximum OD achieved for all three mutants was less than observed for WT, an observation that was most obvious for the double mutant. The discrepancies in maximum growth between WT and mutants was more pronounced at 42°C. These data suggest that *clpP1* and *clpP2* may play roles in the transition to stationary phase and/or spore formation and in temperature stress recovery.

**Figure 2.**
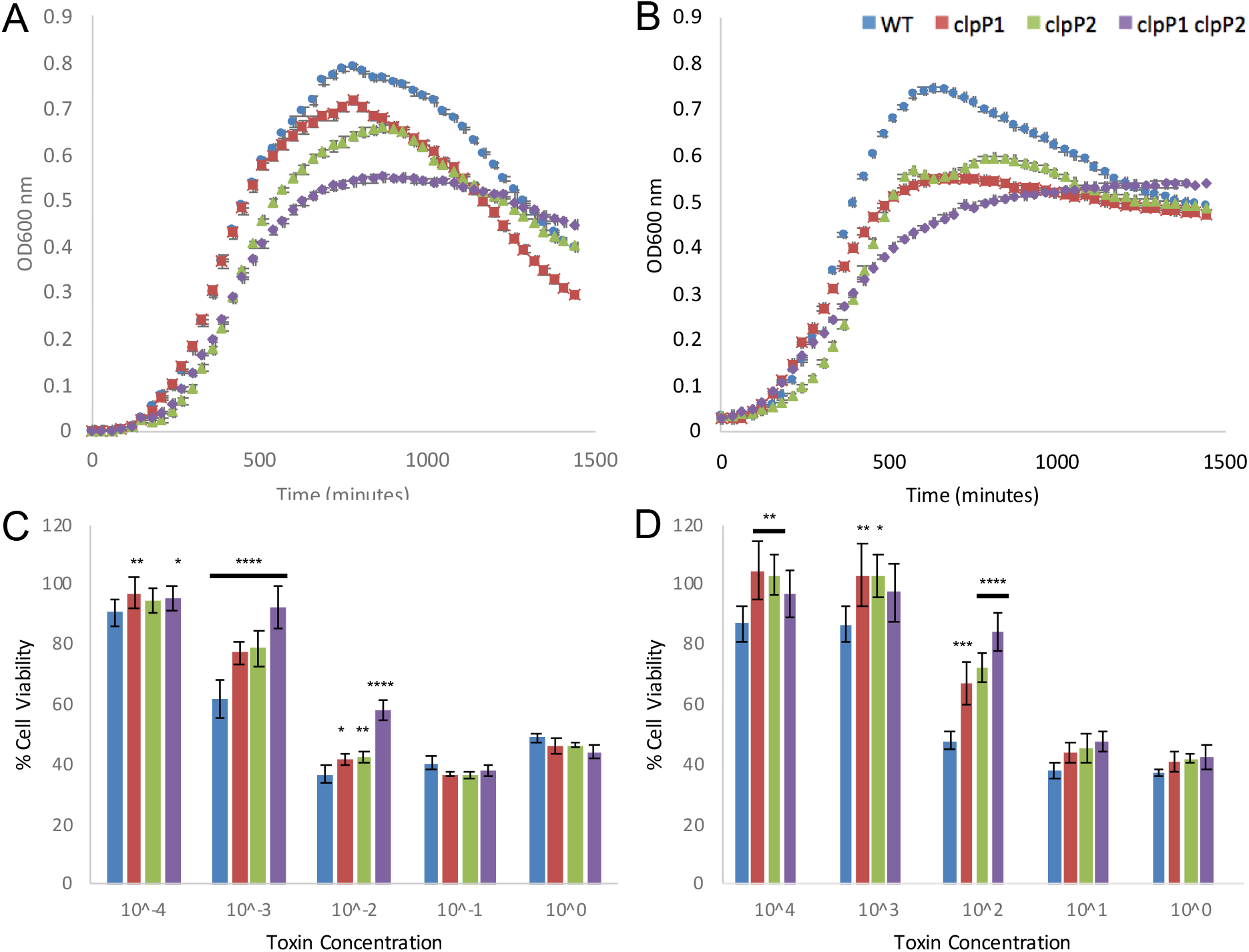
Growth profiles for *clpP* mutants and cytotoxicity TcdB titer assay. *C. difficile* strain 630 WT, *clpP1* mutant, *clpP2* mutant, and *clpP1/clpP2* double mutant were grown anaerobically at 37°C (A) and 42°C (B) and the optical density at 600 nm (OD600) was continually monitored for 25 h. Natively purified TcdB from *C. difficile* 630 was used to titer relative TcdB toxin levels against CHO (C) and Caco-2 (D) mammalian cells. Assays were performed in technical and biological replicate and standard deviation is presented here for each data point.

Research by Baker and colleagues has demonstrated that *Pseudomonas aeruginosa* ClpP1 and ClpP2 maintain separate roles in virulence (Hall, Breidenstein et al. 2017). *P. aeruginosa* ClpP1 maintains an important role in toxin production, while *P. aeruginosa* ClpP2 influences late-stage adhesion and microcolony formation. Each ClpP isoform in *P. aeruginosa* is differentially expressed and maintains specialized functions. We anticipated a similar outcome in *C. difficile*, with the two ClpP isoforms maintaining separate but important roles. Growth data indicates an important role for ClpP1 in heat-shock recovery, while the *clpP2* mutant maintains a consistent growth profile at 37°C and 42°C.

The clinical pathology of CDI is directly tied to the production of two exotoxins, TcdA and TcdB, which glycosylate Rho family GTPases in host cells and disrupt cell morphology, induce cell death, and elicit a strong inflammatory response. To assess the effects of *clpP* mutation on toxin biology, we evaluated the supernatants of WT and mutant cell cultures for the presence and toxicity of the TcdB toxin, one of the major toxins produced by *C. difficile* (Voth and Ballard 2005). In a TcdB toxin titer assay against CHO (**Figure 2C**) and Caco-2 cells (**Figure 2D**) all mutants exhibit one to two-fold lower cytotoxicity in comparison to WT. The *clpP1* and *clpP2* mutants exhibit similarly reduced cytotoxicity, while the *clpP1/clpP2* double mutant demonstrates an additive attenuation. This effect was not cell-specific, which could indicate that each ClpP isoform contributes to toxin production or that either ClpP isoform is sufficient to rescue toxin production.

### *C. difficile* ClpP1 and ClpP2 are both required for heat-stable spore production

Given that the growth data suggested a role for ClpPs in *C. difficile* transition to stationary phase and, potentially, sporulation, we wanted to determine the essentiality of ClpP1 and ClpP2 in *C. difficile* sporulation. To do this, we analyzed the apparent sporulation behavior of *clpP* mutants using phase-contrast microscopy and assessed the heat stability of spores produced by each mutant compared to WT using a Heat Resistance Assay (HRA) described previously (Putnam, Nock et al. 2013, Fimlaid, Jensen et al. 2015, Fimlaid, JEnsen et al. 2015, Edwards and McBride 2016, de la Puebla, Giacalone et al. 2020). Sporulation was visualized by phase contrast microscopy after ~22 h growth on 70:30 solid medium for all data presented in **Figure 3**. Total vegetative cells and phase-bright spore structures from multiple images were counted. Representative images for each strain are presented in **Figure 3A**. The number of phase-bright spores and forespores per image was determined (**Figure 3B**). The percent of phase-bright spores and forespores in the total population (sporulation efficiency) was then calculated for all strains and compared to WT. The sporulation efficiency is defined as [# spores / (# vegetative cells + # spores)] × 100.

**Figure 3.**
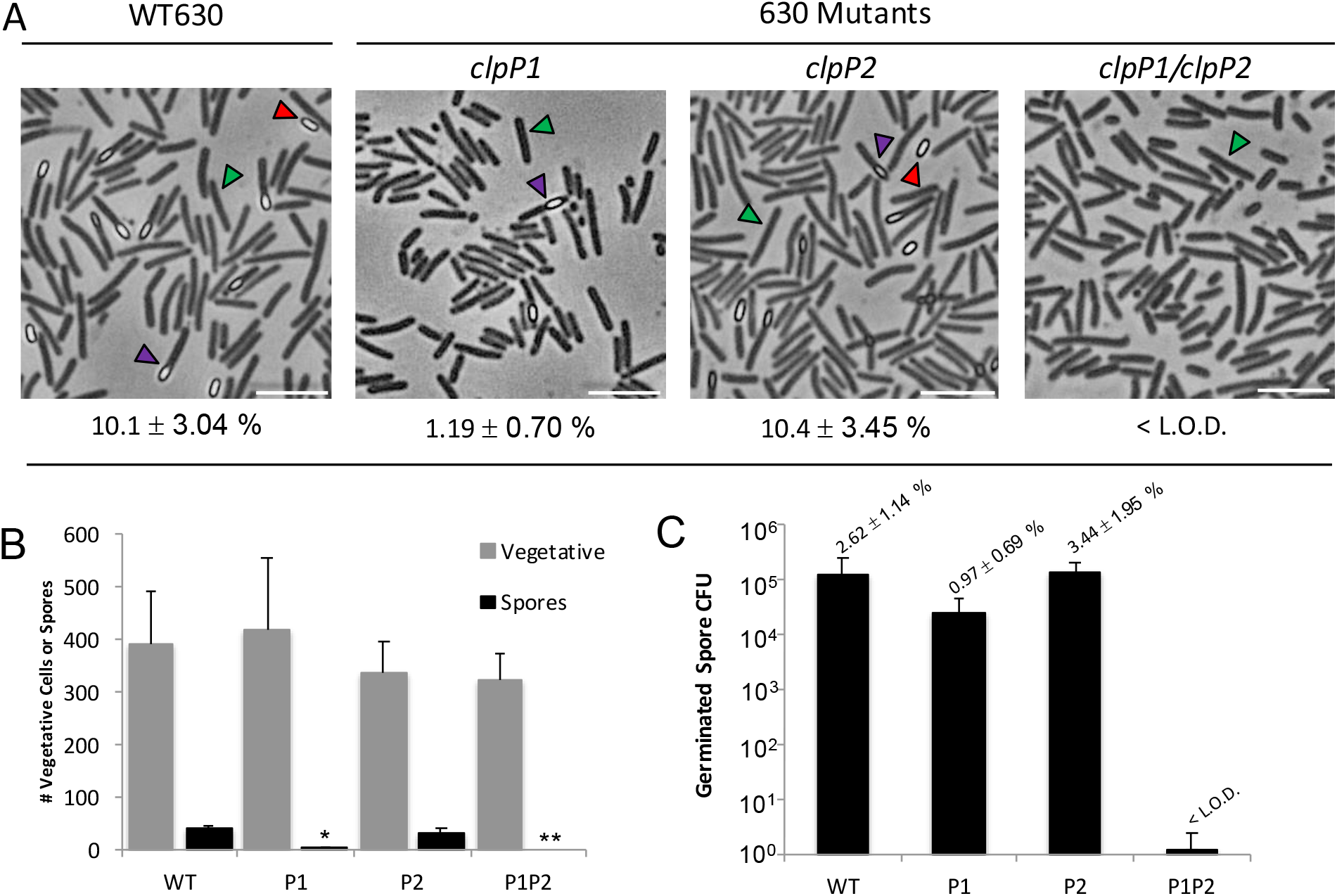
Bright field microscopy and semi-quantitative sporulation rate analyses in *clpP* mutants. (A) Representative phase contrast microscopy images of each strain after 22 h growth on 70:30 plate medium were taken with an Olympus CX41RF fitted with Olympus CCD SC50 and CellSens software (Phase 2, 40x). Vegetative *C. difficile* appear as phase-dark rods (green arrows), while spores appear phase-bright (red arrows). Apparent mother cell containing phase-bright forespore structures are annotated with a purple arrow. White scale bars represent 10 μm. The sporulation efficiency, defined as [# spores / (# vegetative cells + # spores)] × 100, is listed for each strain under the corresponding image (Limit of Detection, L.O.D., 0.001%) (B) Areas were counted under phase-contrast, which show vegetative cells (veg) as phase-dark rods, and spores as small phase-bright ovals either alone or within the mother cell. (C) # of CFU counted per recovery plate post heat-treatment of sporulating cultures (L.O.D., 1 × 10^2^ CFU). The % germination efficiency is presented above each strains bar and is defined as (# heat treated / # CFU untreated) × 100. Experiments were performed in technical and biological triplicate. The means and standard error of the means from three biological replicates are shown with significance determined by two-tailed Student’s t-test (**** P≤0.0001, *** P≤0.001, ** P≤0.01, * P≤0.05).

The *clpP1* mutant produced an average of 4 phase-bright spores/forespores and over 400 vegetative cells as counted in multiple phase-contrast images captured from assays run in biological triplicate (**Figure 3B**). This yielded an apparent sporulation efficiency of 1.19 ± 0.70% for the *clpP1* mutant (**Figure 3A**), which is 13% that of WT. These data suggest that loss of ClpP1 affects the number of spores formed by *C. difficile* but that this isoform is not required for spore production in total. The *clpP1* complement reverts back to WT sporulation, as detected by phase-contrast microscopy, with a sporulation efficiency of 10.8 ± 4.06% (data not shown), comparable to WT sporulation efficiency (10.1 ± 3.04%, **Figure 3A**). Conversely, spore production in the *clpP2* mutant mirrors WT, producing an average of 32 spores per ~400 vegetative cells (**Figure 3B**) and a sporulation efficiency of 10.4 ± 3.45% (**Figure 3A**), 100% that of WT. With this in mind, we expected the *clpP1/clpP2* double mutant to produce spores comparably with the *clpP1* mutant. Much to our surprise, however, no spores were detected via phase-contrast microscopy in biological and technical replicates for the *clpP1/clpP2* double mutant (**Figure 3A**), thus sporulation efficiency for the *clpP1/clpP2* double mutant was determined to be below the assay limit of detection (LOD = 0.001%).

To determine the viability of spores produced by *clpP* mutants compared to WT *C. difficile*, bacterial colonies producing spores, as detected by phase-contrast microscopy (e.g., **Figure 3A**), were harvested from 70:30 plates after ~22 h incubation. Half of the sample was heat-treated, plated and incubated on BHIS plates supplemented with sodium taurocholate to promote germination. The presence of viable, germinating spores was detected after 24 h incubation as CFUs and compared to untreated (i.e., no heat challenge) controls for each strain. This provided us with the germination efficiencies for each strain (**Figure 3C**), defined as (# CFUs heat treated / # CFUs untreated) × 100. The *clpP1* mutant produced 10-fold less CFUs from heat-treated samples compared to WT (**Figure 3B**) and a germination efficiency of 0.97 ± 0.69% (**Figure 3C**), suggesting that in addition to the *clpP1* mutant producing less apparent spores (**Figure 3B**), fewer of these spore structures were mature and/or withstood heat tolerance, a measure of viability. The *clpP1* complement reverted to WT behavior with a germination efficiency 132 ± 29% that of WT (data not shown). As with sporulation efficiency, the viability of heat-treated spores did not vary significantly when the *clpP2* mutant germination efficiency was compared to the WT. The *clpP2* mutant produced an average of 1.33 × 10^5^ CFUs post-heat treatment with a germination efficiency of 3.44 ± 1.95%, similar to that of WT germination efficiency (2.62 ± 1.14%) (**Figure 3C**). Finally, the *clpP1/clpP2* double mutant produced significantly less heat-tolerant spore CFUs compared to WT (**Figure 3C**), producing an average of 1.22 CFUs in undiluted heat-treated samples (**Figure 3C**) across 3 biological replicates with 3 technical replicates per experiment. That the *clpP1/clpP2* double mutant had a few detectable CFUs in the undiluted sample post heat-treatment could be attributed to the production of a minimal number of spores that are undetected by phase-contrast or TEM methods. Regardless, it is clear that a significant defect in sporulation exists for the *clpP1/clpP2* double mutant. To summarize, when spores were purified under anaerobic conditions, the *clpP1* mutant produced less spores than WT and the *clpP1/clpP2* fails to produce spores at levels of detection, while the sporulation profile of the *clpP2* mutant mirrors that of WT. Additionally, the germination efficiency of the *clpP1* mutant and the *clpP1/clpP2* double mutant are decreased compared to WT.

### Spore morphology of *clpP* mutants

The spore morphology of *C. difficile* strain 630 and the *clpP* mutants was characterized by TEM imaging of whole spores (**Figure 4**). The general structure of *C. difficile* spores consists of an outer spore coat, outer membrane between the coat and the abutting layer, the cortex, a germ cell wall, and the inner membrane surrounding the core (Paredes-Sabja, Shen et al. 2014). Herein, spores that maintained at least a core, cortex, and coat were considered “typical” spores and those that lacked any of the three were considered “atypical”. The majority of the spores produced by the *clpP1* and *clpP2* mutants had an obvious inner core, cortex, and an electron dense spore coat, which confirms that both of these mutants are capable of producing free spores that retain WT morphology in regard to those features (**Figure 4A**). The *clpP1* mutant produced 69.6 ± 8.3% typical spores and the *clpP2* mutant produced 75.6 ± 3.2% typical spores, while WT produced 82.4 ± 1.3% typical spores (**Figure 4C**). The *clpP1* mutant samples exhibited double the number of atypical spores compared to WT (30.4 ± 8.3% and 17.6 ± 1.3%, respectively). No typical spores were detected for the *clpP1/clpP2* double mutant (**Figure 4B**), hence 100% of the structures counted in purified spore preps for the double mutant were considered atypical (**Figure 4C**).

**Figure 4.**
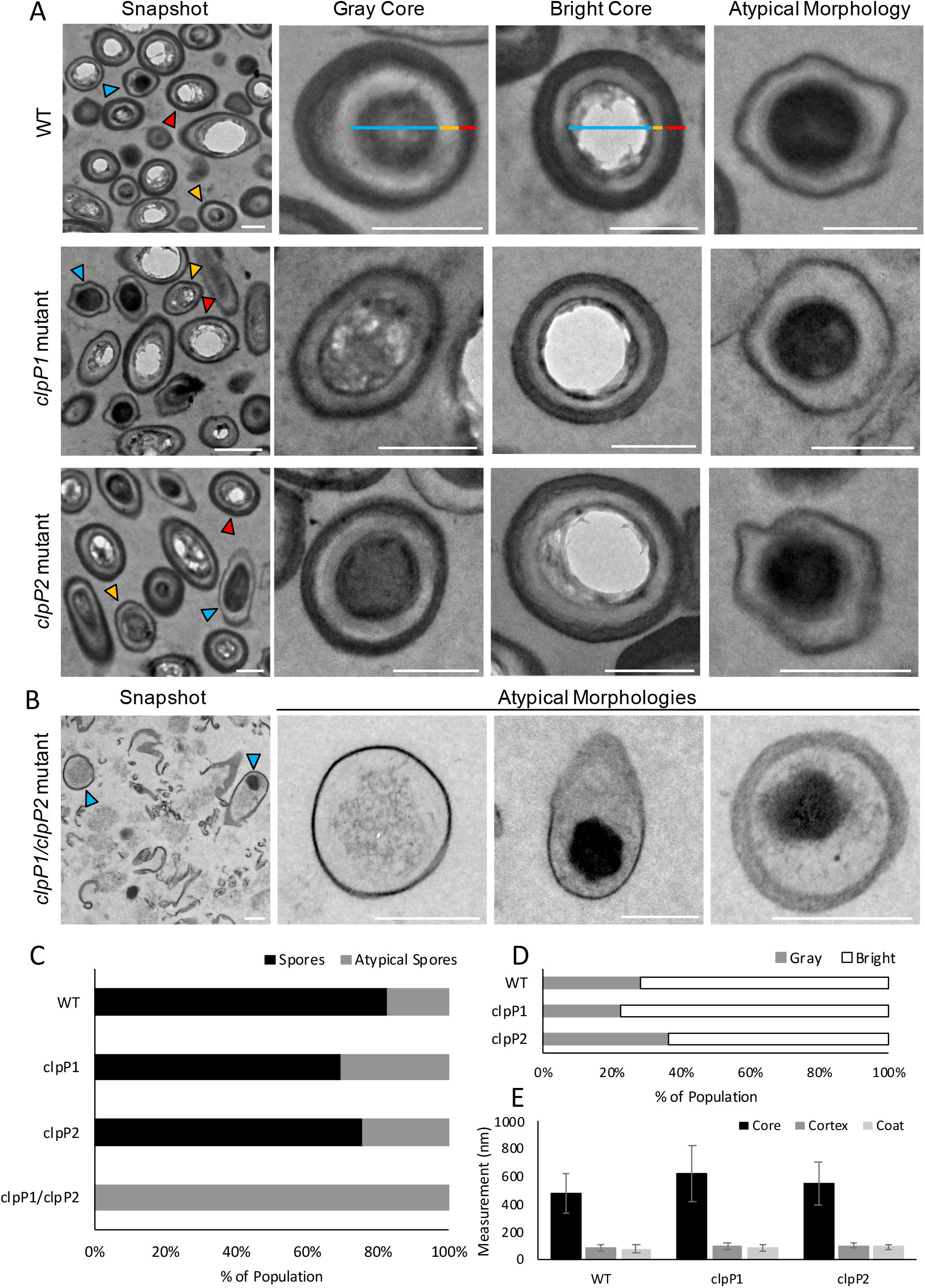
Cellular morphology of each strain in sporulating culture. (A) Representative TEM images of *C. difficile* strain 630 WT, *clpP1* mutant, *clpP2* mutant, and *clpP1/clpP2* double mutants as “Snapshots” of the spore morphologies present in each sample. Representative TEM gray and bright cores (yellow and red arrows, respectively) as well as atypical spore morphologies (blue arrows) are presented for WT, *clpP1* and *clpP2* mutants. (B) Examples of three different atypical spore morphologies as well as a snapshot of the atypical spore community for the *clpP1*/*clpP2* double mutant. Scale bars designate 500 nm for all images. (C) The average number of structures was determined for each sample in triplicate for a minimum of 100 structures per strain. The percentage of total spores (black) and the total percentage of atypical spore morphologies (gray) was determined out of total structures. (D) The percentage of phase-gray spores (gray) was also compared to the percentage of phase-bright spores (white) for WT, *clpP1* and *clpP2* mutants only, as the *clpP1/clpP2* double mutant did not produce typical spore morphologies. The coat proteins and denseness of the core in mature spores often prohibits total resin and stain penetration, causing the core to appear hollow (bright). (E) The average spore core (blue line), cortex (yellow line), and coat (red line) thickness was determined for each sample from 50 total spores for WT, *clpP1* and *clpP2* mutant spores in triplicate (n=150). Example measurement lines are indicated on WT gray and bright spore images. Four total measurements were taken from different angles for each spore and the total measurement (nm) was averaged.

For the typical spores examined in WT, *clpP1* mutant, and *clpP2* mutant samples, the core was either electron dense and appeared dark/gray or the spores appeared bright (**Figure 4A**). While the denseness of the coat in mature spores often prohibits total resin and stain penetration, causing the core to appear hollow or bright, this can also be due to a lack of electron density altogether. Of note is that the *clpP1* mutant samples contained more spores with bright cores compared to WT and the *clpP2* mutant (**Figure 4D**). The *clpP1* mutant produced slightly less gray/dark cored spores (22.3 ± 8.5% of total) than WT (28.3 ± 6.2% of total), whereas the *clpP2* mutant produced more gray/dark cored spores (36.3 ± 5.7% of total) than WT (**Figure 4D**). As can be seen in the representative images (**Figure 4A**), it is likely that the bright spore cores are due to incomplete stain penetration rather than an absence of electron density entirely since edges of core material and areas of density (darkness) running through bright cores can be seen (see “Snapshot” images, **Figure 4A**). The interest in these differing values lies in the correlation between TEM analysis and the sporulation efficiency data presented earlier in this work, which demonstrates a diversion from WT sporulation for the *clpP1* mutant and loss of sporulation in the *clpP1/clpP2* double mutant. TEM data further support the evidence that combined loss of function of ClpP1 and ClpP2 results in a strain that can no longer form spores. It is tempting to speculate that the propensity of the *clpP1* mutant to yield less spores compared to WT (**Figure 3B**) and the production of majority phase-bright spore cores for the *clpP1* mutant provides some congruence between data. However, more studies are clearly needed to determine the exact role that ClpP1 plays in the production of mature spores. It will be interesting, for instance, to determine in future work whether loss of ClpP1 causes over-abundance of late phase sporulation proteins under sporulating conditions, including many of the spore coat proteins, thus yielding spores that are meretricious and impenetrable to the TEM stains. This could cause *C. difficile* to make fewer spores due to improper allocation of resources as well as the production of spores that do not germinate as efficiently as WT.

To determine if there were differences in core, cortex, and/or spore coat dimensions between WT and the mutants that produced free spores, measurements of these structures were performed (**Figure 4E**). On average, the *clpP1* and *clpP2* mutant spores had a wider cortex and spore coat than WT. The *clpP1* mutant cortex measured 100 ± 21.3 nm and the coat measured 88.5 ± 21.8 nm. The *clpP2* mutant cortex measured 101 ± 19.1 nm and the coat measured 93.9 ± 19.7 nm, while the WT cortex measured 84.6 ± 22.2 nm and the coat measured 78.8 ± 28.2 nm. Additionally, the *clpP1* mutant had the widest core measurement (623 ± 203 nm) compared to WT (480 ± 140 nm) and the *clpP2* mutant (552 ± 151 nm). However, none of these size differences were statistically significant. For WT and the *clpP1* and *clpP2* mutants the percentage of spore structure components over the total spore diameter were similar for core, cortex, and coat (73-75%, 12-14%, and 12-13%, respectively) (Supplemental Table S2). That there was no strong correlation between spore dimensions and behavior has been observed previously for *Clostridium sordellii* (Rabi, Turnbull et al. 2017). Additionally, though the sporulation process is tightly regulated, it has been proposed previously that variations in spore structure might exist as an advantage to bacterial survival (Faille, Lequette et al. 2010). The greatest difference between strains was observed for the *clpP1/clpP2* double mutant, whereby no typical spores were isolated and thus available for visualization by TEM. Another notable difference between WT and *clpP* mutants lies in the number of atypical spores between the *clpP1* mutant (30.4 ± 8.3%) and *clpP2* mutant (24.4 ± 3.2%) compared to WT (17.6 ± 1.3%). The *clpP* mutants ultimately produce more atypical spores than WT.

### Comparative proteome profiling of ClpP Mutant Strains

To understand the influence genetic manipulation of *clpP1* and *clpP2* has on the *C. difficile* 630 proteome, the global protein abundance levels of the *clpP* mutants were compared to WT cells in five biological replicates. Reproducibility among biological replicates and the differences between WT and *clpP* mutants was assessed using a principal component analysis (PCA) (**Figure 5A**). The PCA showed high reproducibility among replicates based on the close clustering of samples for each of the strains examined with the exception of a single outlier for the *clpP2* mutant. The PCA also showed the close clustering of *clpP1* and *clpP2* mutants and the distinction of these mutants from WT and the *clpP1/clpP2* double mutant. Not unexpectedly, the *clpP1/clpP2* double mutant appears distinct from WT and the individual *clpP* mutants. The *clpP1* and *clpP2* samples cluster closely together, suggesting that the phenotypical differences in sporulation between these two mutants may not be due to global proteomic differences, but rather are reflective of differences in expression of very specific proteins. Heat map clustering of the total unique identified proteins further demonstrates the differential abundance in each group (**Figure 5B**) and reveals the portions of the proteome, as exemplified in **Figure 5C**, that show significant diversion from WT protein abundance for certain portions of the proteome. In order to investigate this further, we independently compared the *clpP* mutants to WT.

**Figure 5.**
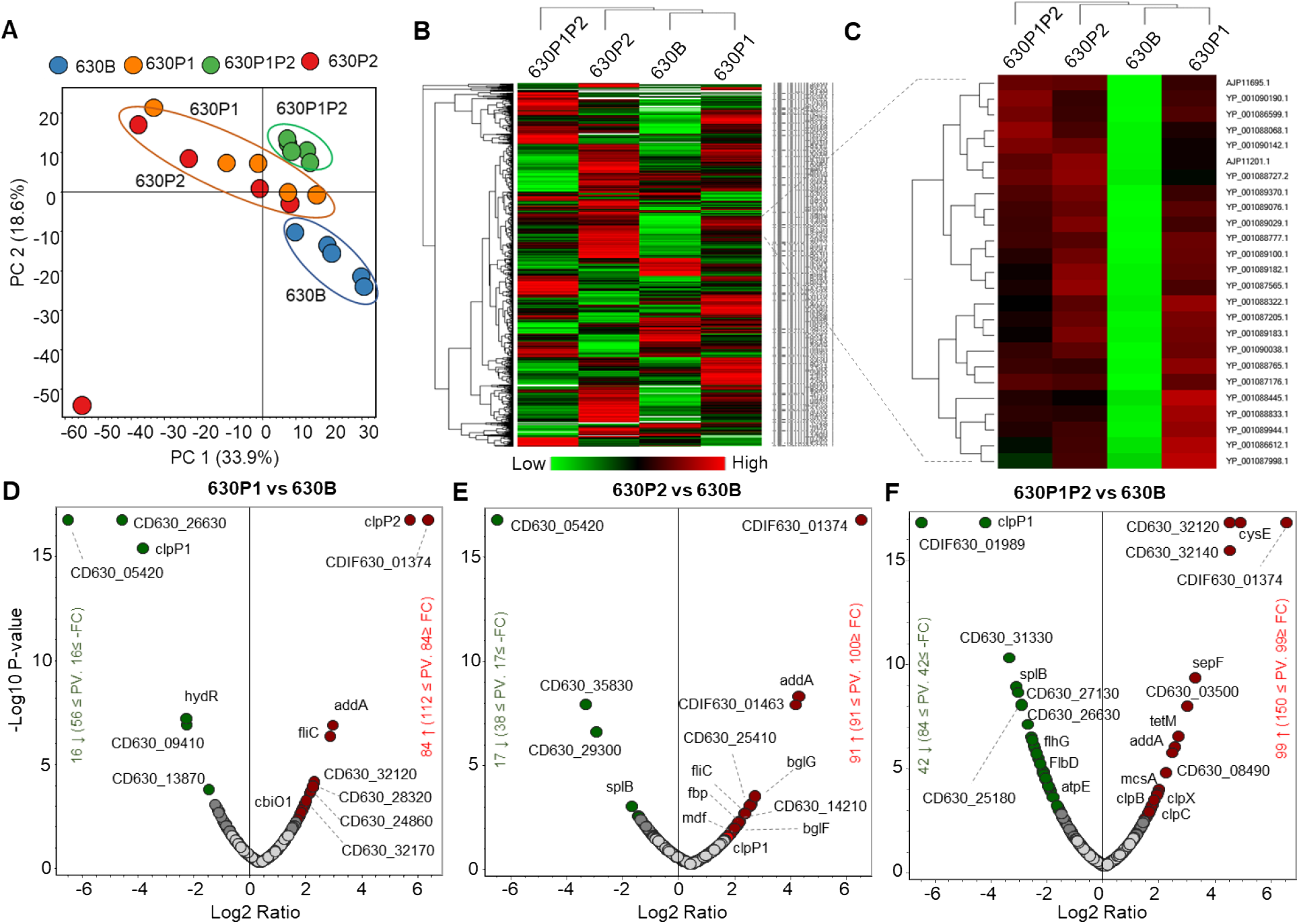
Comparative label-free quantitative proteomic analysis of *C. difficile* strain 630 (WT) and mutant strains for *clpP1* and *clpP2* mutants, and *clpP1/clpP2* double mutant. Total proteome identified from five biological replicates were subjected to quantitative analysis. (A) Principal component analysis (PCA) shows close clustering of the total normalized protein abundance (peak area) of the replicates of *clpP1* and *clpP2* mutant strains, however distinct from WT and *clpP1/clpP2* double mutant strains. (B-C) Heat map clustering of the total 1730 identified and quantified unique proteins shows differential abundance in each group. (C) Example of a group of proteins showing very similar expression pattern in all three mutant strains comparative to WT. (D-F) Volcano plot analysis of the 1730 proteins subjected for quantitative analysis. Significant increases and decreases in abundance of proteins in *clpP1* mutant vs WT (D), *clpP2* mutant vs WT (E), and *clpP1/clpP2* double mutant vs WT (F) are marked as red and green, respectively. X-axis represents the protein abundance log2 ratio and Y-axis represents the statistically significance with –log10 of p-value. The gray circles are non-significant (p>0.05) and below the threshold protein abundance fold change (1.5-fold).

Label-free quantitative proteomic analysis shows, the *clpP1* mutant exhibits 84 significantly up- and 16 significantly down-regulated proteins (**Figure 5D**), the *clpP2* mutant reveals 91 significantly up- and 17 significantly down-regulated proteins (**Figure 5E**), and the *clpP1/clpP2* double mutant exhibits 99 significantly up-regulated and 42 significantly down-regulated proteins (**Figure 5F**) when compared with the WT. While proteins impacted by loss of ClpP function have been investigated in other organisms (Weichart, Querfurth et al. 2003, Kock, Gerth et al. 2004, Feng, Michalik et al. 2013, Zheng, Wu et al. 2020, Kirsch, Fetzer et al. 2021), this is the first data set demonstrates the impact of ClpP absence on *C. difficile*.

### Comparison of *clpP* mutant with Spo0A-regulated and mature spore proteomes

Researchers have examined the *C. difficile* proteome previously to understand heat stress response (Jain, Graham et al. 2011), examine the differences between *in vitro* and host environment (Janvilisri, Scaria et al. 2012), compare historic and hypervirulent proteomes (Chen, Scaria et al. 2013), generate a large inventory of *C. difficile* proteins on different standard lab growth media (Otto, Maass et al. 2016), and generate proteomic signatures of drug-stressed *C. difficile* (Maass, Otto et al. 2018). A comparison of our work to previous research demonstrates the breadth of proteins identified in this study (**Figure 6A**). For a better understanding of the role ClpP plays at the protein-level in sporulation, we further compared our proteome dataset with the Spo0A-regulated proteome (Pettit, Browne et al. 2014) and the mature spore proteome (Lawley, Clare et al. 2009), including only the proteins that were significantly up- or down-regulated (p-value < 0.005). These proteome data sets were chosen for comparison due to their direct relationship to sporulation. Spo0A is an early-stage sporulation transcription factor that is often called “the master spore regulator” due to its central role in initiation of the sporulation cascade (Hoch 1993, Pettit, Browne et al. 2014). Comparing our data to the proteins up- or down-regulated in the absence of Spo0A (Pettit, Browne et al. 2014) offers a robust tool for understanding similarities between an asporogenic strain and our *clpP* mutants. In addition to visualizing our data in tandem with the Spo0A-regulated proteome, we aimed to understand the proteins present in the *clpP* mutants compared to those identified in the highly purified mature spore proteome (Lawley, Clare et al. 2009). We captured 64.5% of these previously reported spore proteomes with 427 of our 1,730 total identified proteins represented in these studies (**Figure 6B**).

**Figure 6.**
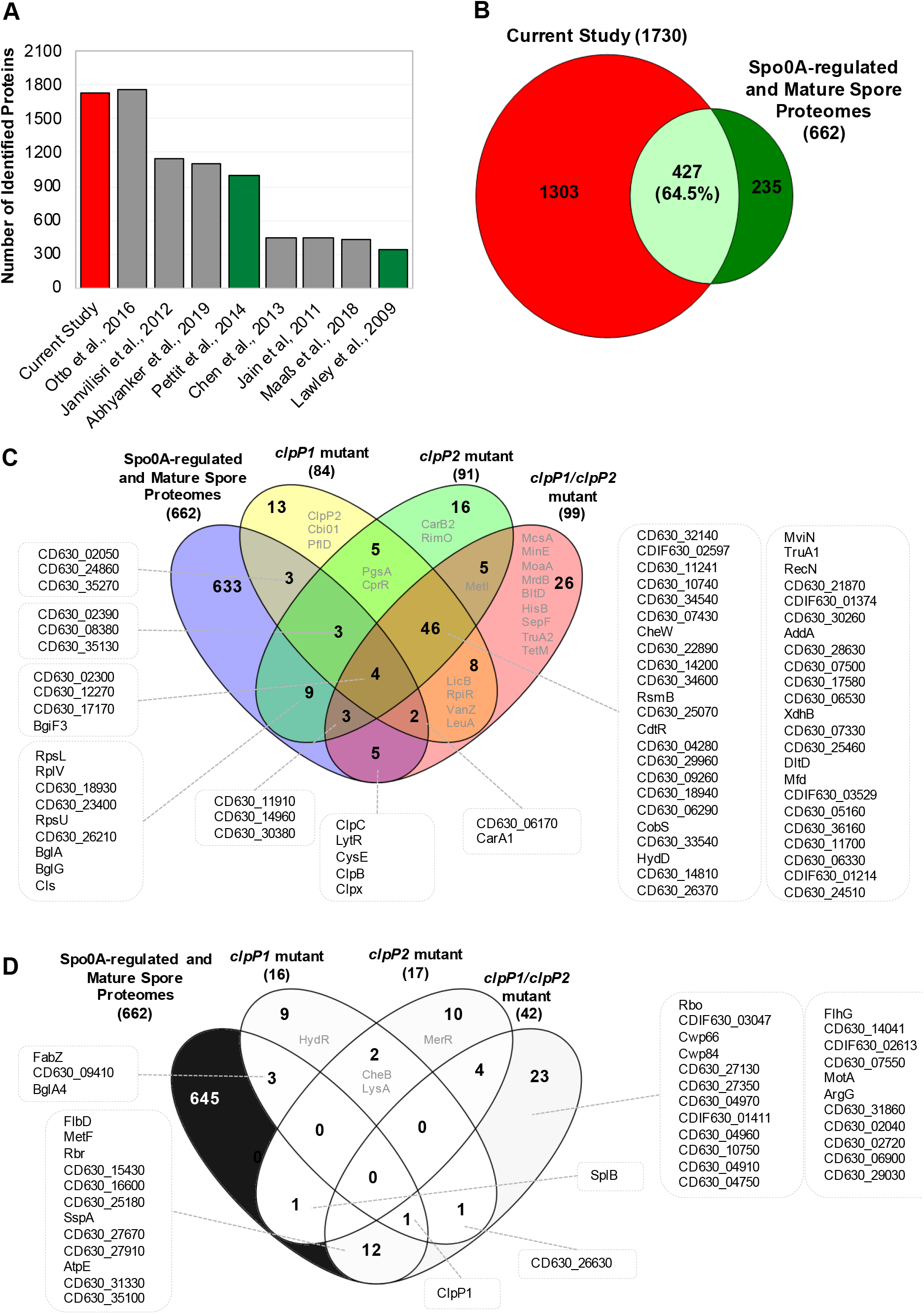
Comparative *Clostridioides difficile* proteomics. Data used to generate this figure can be found in Supplemental Table S4. (A) Bar diagram represents the number of proteins identified from different proteomics studies of *C. difficile*. Studies used in comparative analysis (B-D) are indicated with green bars. (B) Proportional Venn diagram analysis shows the spore associated protein coverage of the current study with the Spo0A-regulated (Pettit, Browne et al. 2014) and mature spore proteomes (Lawley, Clare et al. 2009). C and D represent the list of unique and overlapped significantly up-regulated (C) and down regulated (D) proteins of in comparison with the Spo0A-regulated and mature spore proteome datasets.

The comparison between the significantly up-regulated proteins in the *clpP* mutants vs Spo0A-regulated and mature spore proteomes (662 total proteins) is presented in **Figure 6C**. As expected, a group of unique protein sets were identified in each ClpP mutant strain compared with the spore proteome data set. For instance, the *clpP1* mutant exhibits 13 unique proteins, which includes ABC transporters (CD630_01000, CD630_35250) and a putative dehydratase (PflD). We identified 16 proteins unique to the *clpP2* mutant including proteins involved in the PTS system (CD630_10760), transport (CD630_21720), and pyrimidine metabolism (CarB2). Similarly, the *clpP1/clpP2* mutant proteome reveals 26 unique proteins including proteins involved in arginine and proline metabolism (BltD), histidine metabolism (HisB), and folate biosynthesis (MoaA). All of the *clpP* mutants share a total of 46 common up-regulated proteins, including two-component systems (CheW, CdtR, DltD, and CD630_26370) and proteins involved in metabolic pathways (e.g. fructose and mannose metabolism, CD630_10740, and starch and sucrose metabolism, CD630_25460). The *clpP2* mutant shares 9 common proteins with the *spo0A* mutant and the mature spore proteome, including ribosomal proteins (RpsL, RpsU, and RplV) and bgl system proteins (BglA and BglG). Proteins identified in *clpP1* mutant and proteomes related to sporulation were few and primarily involved in transport (CD630_24860 and CD630_35270). FliC up-regulation was common between the single *clpP* mutants and the up-regulated proteins in the *spo0A* mutant proteome and the mature spore proteome. Whereas the overlap between the *clpP1/clpP2* double mutant and the comparison proteomes totaled 5 proteins including known ClpP co-chaperones ClpC (CD630_00260) and ClpX (CD630_33040).

Similarly, the significantly down-regulated proteins of *clpP* mutants were also compared with the Spo0A-regulated, mature spore proteome (**Figure 6D**). A total of nine proteins were unique to the *clpP1* mutant included 2 putative TetR family transcriptional regulators (HydR, CD630_25270). We found that the *clpP2* mutant had 10 unique down-regulated proteins including a transcriptional regulator from the MerR family and an ABC transporter-like protein (CD630_31950). The *clpP1* mutant and the *clpP2* mutant share two common down-regulated proteins (CheB and LysA). Few other similarities in down-regulated proteins are seen when comparing the *clpP* mutants with each other. For instance, a putative signaling protein (CD630_26630) shared between the *clpP1* mutant and *clpP1/clpP2* double mutant and type IV pilus assembly protein (CD630_35120) and putative nucleases (CD_00560 and CDIF630_01989) shared between the *clpP2* mutant and the *clpP1/clpP2* double mutant. A total of 23 down-regulated proteins were unique to the *clpP1/clpP2* mutant when compared with the spore proteome data set. These proteins included cell wall (CD630_27130, CD630_27350) and cell surface (Cwp66 and Cwp84) proteins as well as proteins involved in flagellar assembly (FlhG and MotA) and a predicted oxygen-stress response protein (Rbo). The *clpP1* mutant shares few proteins with the proteomes used for comparison, including proteins involved in fatty acid biosynthesis (FabZ) and glycolysis (BglA). The *clpP1/clpP2* double mutant exhibits 12 down-regulated proteins in common with the *spo0A* mutant and mature spore proteomes. These similarities include cell surface (CD630_25180 and CD630_27670), cell wall binding (CD630_27910) and flagellar (FlbD) proteins as well as a small, acid-soluble spore protein (SspA).

These comparative analyses provide valuable information of potential ClpP proteolytic targets for future studies. For example, the *clpP1/clpP2* double mutant, which has a severe sporulation defect, is the only strain in this comparison to have downregulated Cwp66 and Cwp84, both of which are expressed during host infection with *C. difficile* suggesting a role for these proteins in both sporulation (this study, (Abhyankar, Zheng et al. 2019)) and virulence (Pechine, Janoir et al. 2005). Additionally, insight was gained into the similarities between the *clpP* mutant proteomes and how they diverge from WT. For instance, the *clpP1* mutant exhibits a 50-fold change in abundance of ClpP2 compared to WT and the other mutants, while no increase in ClpP1 is obvious when ClpP2 is absent. This data further supports our previous claim that the ClpP1 and ClpP2 isoforms of *C. difficile* are capable of functioning in an uncoupled fashion. This also suggests that ClpP2 may serve a rescue function in the absence of ClpP1 (Lavey, Shadid et al. 2019). While these data are intriguing, we wanted to examine proteins specifically involved in sporulation by separating the proteome data sets used for comparisons in **Figure 6**, particularly because one data set demonstrates proteins that are up- or down-regulated in the absence of Spo0A and sporulation (Pettit, Browne et al. 2014) and the other indicates proteins that are present in the isolated mature spore proteome (Lawley, Clare et al. 2009). Initially combining these data sets provides an efficient method to obtain an overview of the potential ClpP sporulation protein profile.

### Potential role of ClpP involvement in sporulation

Previous works have integrated data sets from other sporulating pathogens, namely *B. subtilis*, and combined this data with existing knowledge of *C. difficile* sporulation in order to provide insight into the phases of sporulation in *C. difficile* (Fimlaid, Bond et al. 2013, Pettit, Browne et al. 2014, Zhu, Sorg et al. 2018, Shen 2020). Comparison of our data to previous work is shown in **Figure 7**. Both the *spo0A* mutant and the *clpP1/clpP2* double mutant fail to produce viable spores. Proteins that are down-regulated in both the *spo0A* mutant and the *clpP1/clpP2* double mutant are the master spore regulator Spo0A, sporulation-specific sigma factors (SigA2 and SigF), and Stage V sporulation proteins (SpoVS and SpoVG) (**Figure 7**). These protein signatures are shared by the *clpP1* mutant with the exception of SigF, which is up regulated. While this data at first seems compelling, the *clpP2* mutant shares the same down-regulated proteins mentioned above, though maintains typical WT sporulation phenotypes. When mining the data for discrepancies between typical and atypical sporulating strain proteomes SpoIIE was up regulated in the *clpP1/clpP2* double mutant only. This is interesting in that SpoIIE is predicted to be a direct target of Spo0A and, if similar to *B. subtilis*, could play a role in activation of SigF and/or asymmetric septation (Barák and Youngman 1996); however, with a low abundance of SigF and Spo0A in the *clpP1/clpP2* mutant it is presumed that SpoIIE would not have a clear role and is incapable of continuing the sporulation cascade without the other two proteins.

**Figure 7.**
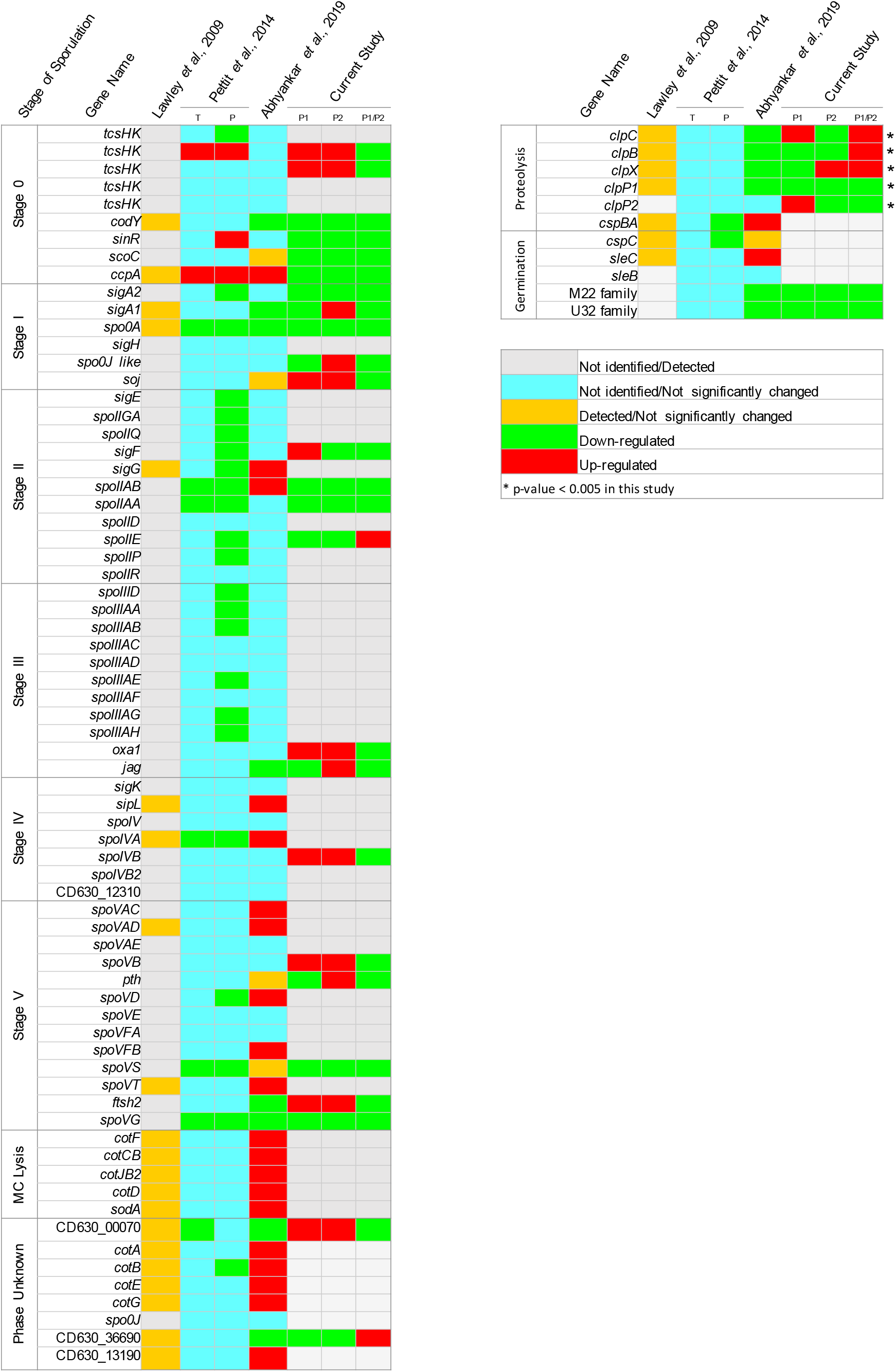
Comparison of known sporulation associated protein expression pattern of *clpP* mutants to the mature spore proteome (Lawley, Clare et al. 2009), *spo0A* mutant proteome and transcriptome (Pettit, Browne et al. 2014), and one-pot spore/vegetative proteome (Abhyankar, Zheng et al. 2019). Gene names for proteins that were “Not Identified/Detected” (gray boxes) were not identified in this study or in the mature spore proteome. Gene names for proteins “Not identified/Not significantly changed” (aqua boxes) were not listed in supplemental tables for the data sets used for comparison. Gene names for proteins that were detected but were either listed as single rep (Abhyankar, Zheng et al. 2019) or the statistical data was unavailable (Lawley, Clare et al. 2009) are represented with yellow boxes. Green and red boxes for other studies (Pettit, Browne et al. 2014, Abhyankar, Zheng et al. 2019) indicate significantly up- (red) or down- (green) regulated proteins. In this study, up- and down-regulated proteins are indicated with red and green boxes, respectively, as well. Only those proteins that were significantly changed (p-value < 0.005) are marked with an asterisk. A more extensive comparison list as well as relevant references can be found in Supplemental Table S5.

Proteins detected in the *clpP1/clpP2* double mutant that were not detected in previous studies but may play a role in sporulation include a two-component system histidine kinase (tcsHK CD630_24920), Soj, Oxa1, SpoIVB, and SpoVB. In fact, all of the identified sporulation proteins in the *clpP1/clpP2* mutant were downregulated with the exception of SpoIIE and an unidentified protein (CD630_36690). The *clpP1* mutant demonstrates a similar protein profile, with most sporulation proteins identified as downregulated with the exception of Soj, SigF, Oxa1, SpoIVB, SpoVB, Ftsh2, and a protein predicted to be involved in the lytic phase of sporulation (CD630_00070). The latter of which is downregulated in the *spo0A* mutant. The *clpP2* mutant however, shows more upregulated sporulation proteins and diverges more frequently from the protein profile of the *spo0A* mutant, which supports the *clpP2* mutant’s ability to produce heat-stable spores.

Because the *clpP1* mutant and the *clpP2* mutant are able to form spores based on phase-contrast microscopy and TEM analysis, even with the lower abundance of Spo0A in these mutants, we focused our attention on differences in protein abundance for these mutants. The tcsHK (CD630_15790) is upregulated in both the *spo0A* mutant and the *clpP1* and *clpP2* mutants. Another tcsHK (CD630_24920) shares the same abundance profile in the *clpP* mutants as CD630_15790. Since the *clpP1* mutant and the *clpP2* mutant both form spores and the *spo0A* mutant did not, these tcsHKs may play a more versatile role in sporulation than spore-specific regulatory proteins like Spo0A. Other regulatory and sensory proteins predicted to be involved in sporulation (CodY, SinR, ScoC, CcpA) are downregulated compared to WT in all *clpP* mutants. Only two of these proteins were found in the mature spore proteome, CodY and CcpA (Lawley, Clare et al. 2009) and only CcpA was upregulated in the one-pot spore/vegetative cell analysis performed recently (Abhyankar, Zheng et al. 2019).

Looking further down the sporulation cascade, we can begin to see greater differences between strains that form spores and those that do not. For instance, *clpP1* and *clpP2* have more upregulated proteins compared to the *clpP1/clpP2* mutant in Stages III-V of sporulation. Jag (SpoIIIJ-associated protein in *B.* subtilis, (Errington, Appleby et al. 1992)) and Pth (SpoVC in *B. subtilis*, (Pettit, Browne et al. 2014)) are upregulated in the *clpP2* mutant and downregulated in the *clpP1* mutant and the *clpP1/clpP2* double mutant, which could suggest these proteins play a significant role in viable spore formation. It is possible that the inability of the *clpP1* mutant to form many heat-stable spores may be due to germination defects, however, we did not capture all of the putative germination proteins and, the ones we did identify, were downregulated in all *clpP* mutant strains and share this expression profile with the one-pot spore/vegetative cell analysis performed previously (Abhyankar, Zheng et al. 2019). Therefore, we cannot form a substantial conclusion in this regard. We also did not identify many of the proteins involved in late-stage sporulation and mature spore release from the mother cell (MC Lysis, **Figure 7**). This is most likely due to sampling methods. Our future work focuses on proteomic analysis of samples from sporulating and germinating cultures as well as the BHIS-grown cells examined in this study.

Finally, in examining the ClpP-specific protein profiles in this study, we found that the ClpP co-chaperones (ClpC and ClpX) are upregulated in the *clpP1/clpP2* double mutant. This suggests that proteins are being tagged for degradation, signaling the production and/or recruitment of co-chaperones to the tagged protein, and then these co-chaperones bound to ClpP degradation targets have nowhere to go if ClpP is missing, thus backing up the system and creating a 3- to almost 4-fold increase in abundance of these co-chaperones in *clpP1/clpP2* double mutants (**Figure 7**). Furthermore, ClpX is more abundant in the *clpP2* mutant and ClpC is more abundant in the *clpP1* mutant potentially signifying preferred co-chaperones (e.g. ClpC for ClpP1 and ClpX for ClpP2).

### Conclusion

Taken together, these results indicate that loss of both ClpP isoforms in *C. difficile* strain 630 generates an asporogenic phenotype. The *clpP1* mutant produces significantly fewer free spores, the *clpP2* mutant produces mature spores similarly to WT, while the *clpP1/clpP2* mutant exhibits multiple phenotypic defects. Based on these data, we speculate that ClpP1 is primarily responsible for proteolytic involvement in sporulation, while ClpP2 behaves more as a rescue protein and the loss of both results in severe sporulation defects. Comparative proteomics revealed that ClpP2 is upregulated in the absence of ClpP1. In contrast, no significant changes were observed for ClpP1 production in the *clpP2* mutant. These collective data suggest even more granular differentiated roles for each ClpP isoform at the initiation of engulfment and for completing engulfment to produce free spores, as is indicated by the defective mutants that produce elongated cells with spore-like structures though generating less heat-resistance colonies. These impacts of ClpP impairment on sporulation as well as the decreased cytotoxicity of *clpP* mutants guide and motivate our future studies into the impact of the loss of ClpPs on the *C. difficile* sporulation biology and the promise of small molecule inhibition of ClpP in *C. difficile* mitigation.

## Materials and Methods

### Strains, media, and growth conditions

All strains are listed in Table 1. All *C. difficile* strains are derivatives of clinical isolate 630. Strains were routinely grown in brain heart infusion (BHI) medium (37 g L^−1^, Millipore Sigma) supplemented with 5 g L^−1^ yeast extract (ThermoFisher Scientific) (Sorg and Dineen 2009). Taurocholate (TA, 0.1% wt/vol, Millipore Sigma), thiamphenicol (TAP, 15 μg mL^−1^, Millipore Sigma), and/or kanamycin (KAN, 50 μg mL^−1^, Millipore Sigma) were supplemented in BHIS medium as indicated. To investigate *C. difficile* sporulation phenotypes, 70:30 medium (70% SMC medium and 30% BHIS medium) (Edwards and McBride 2017) was used in addition to BHIS medium. 70:30 medium contained (per L) 63 g Bacto peptone (Gibco), 3.5 g protease peptone (ThermoFisher Scientific), 11.1 g BHI, 1.5 g yeast extract, 1.06 g Tris base (VWR), 0.7 g (NH_4_)_2_SO_4_ (Millipore Sigma), and 15 g agar (Millipore Sigma). Additives to solid media for sporulation studies include TA (0.1% wt/vol) and TAP (15 μg mL^−1)^ as indicated. *C. difficile* was grown at 37°C in an anaerobic chamber (COY) with headspace of 5-8% CO_2_, 2.5-3.9% H_2_, 0 ppm O_2_, and <32% humidity unless otherwise indicated. Starter cultures were grown on solid medium (typically BHIS-TA) before conducting experiments.

**Table 1.**
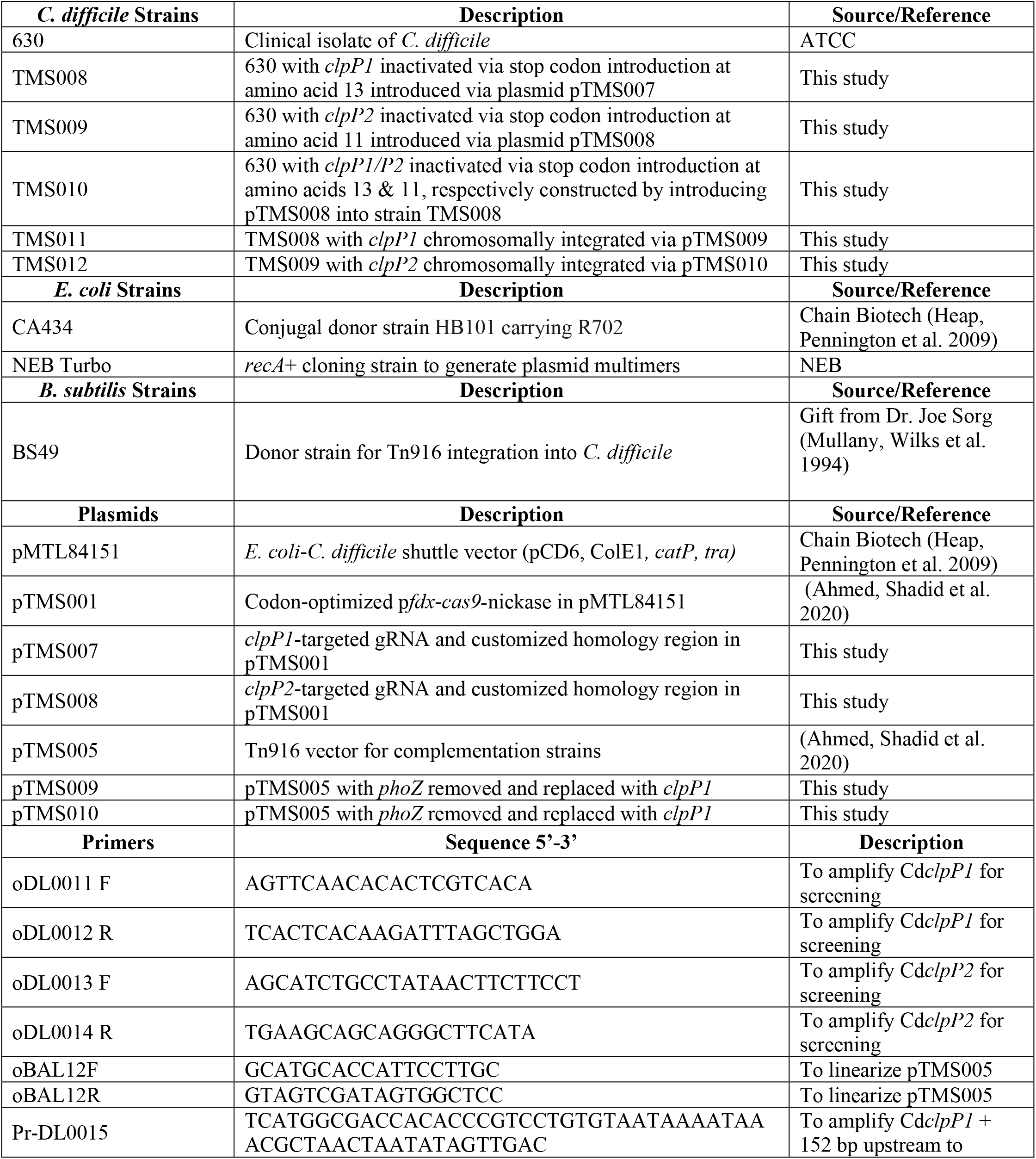
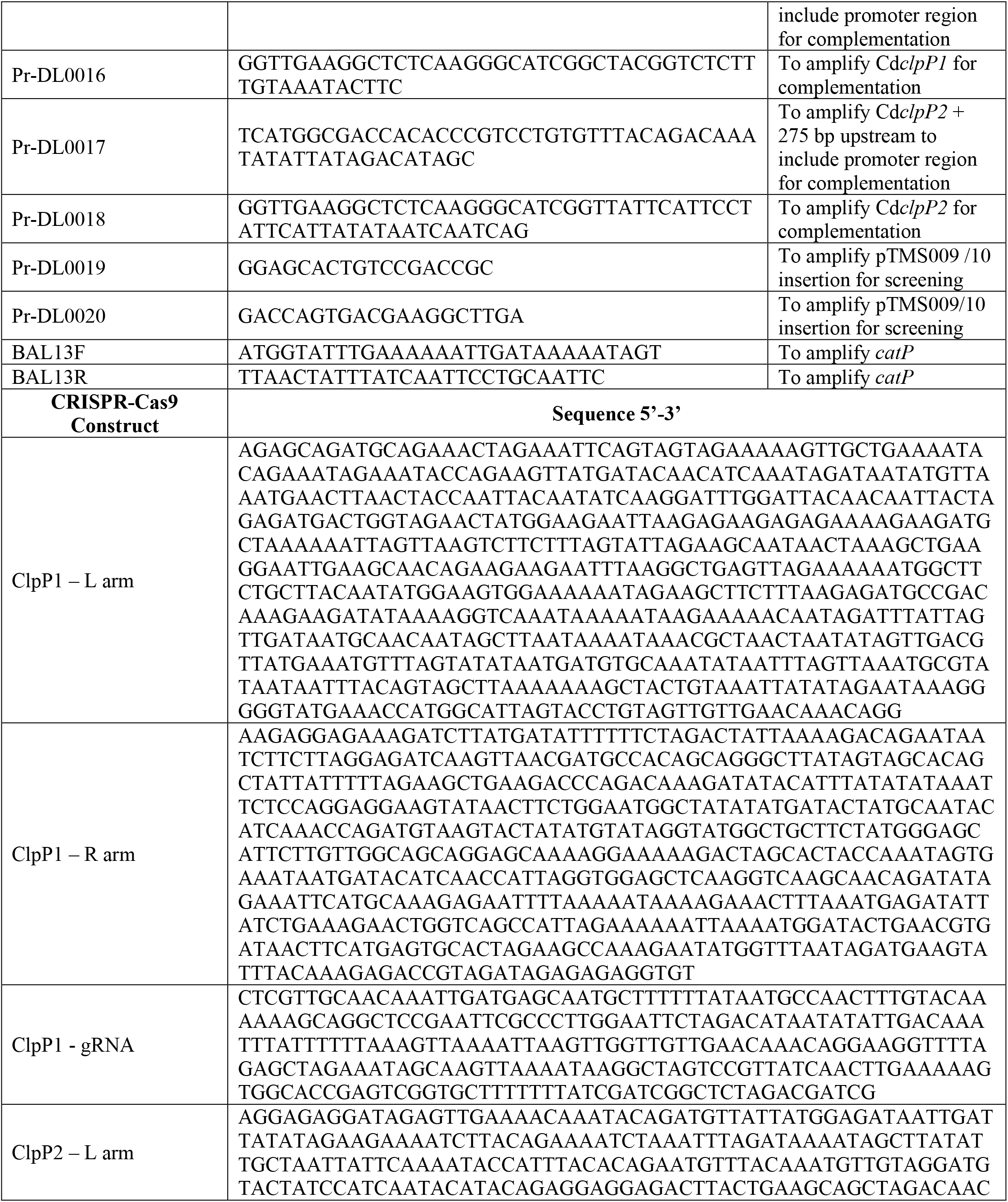
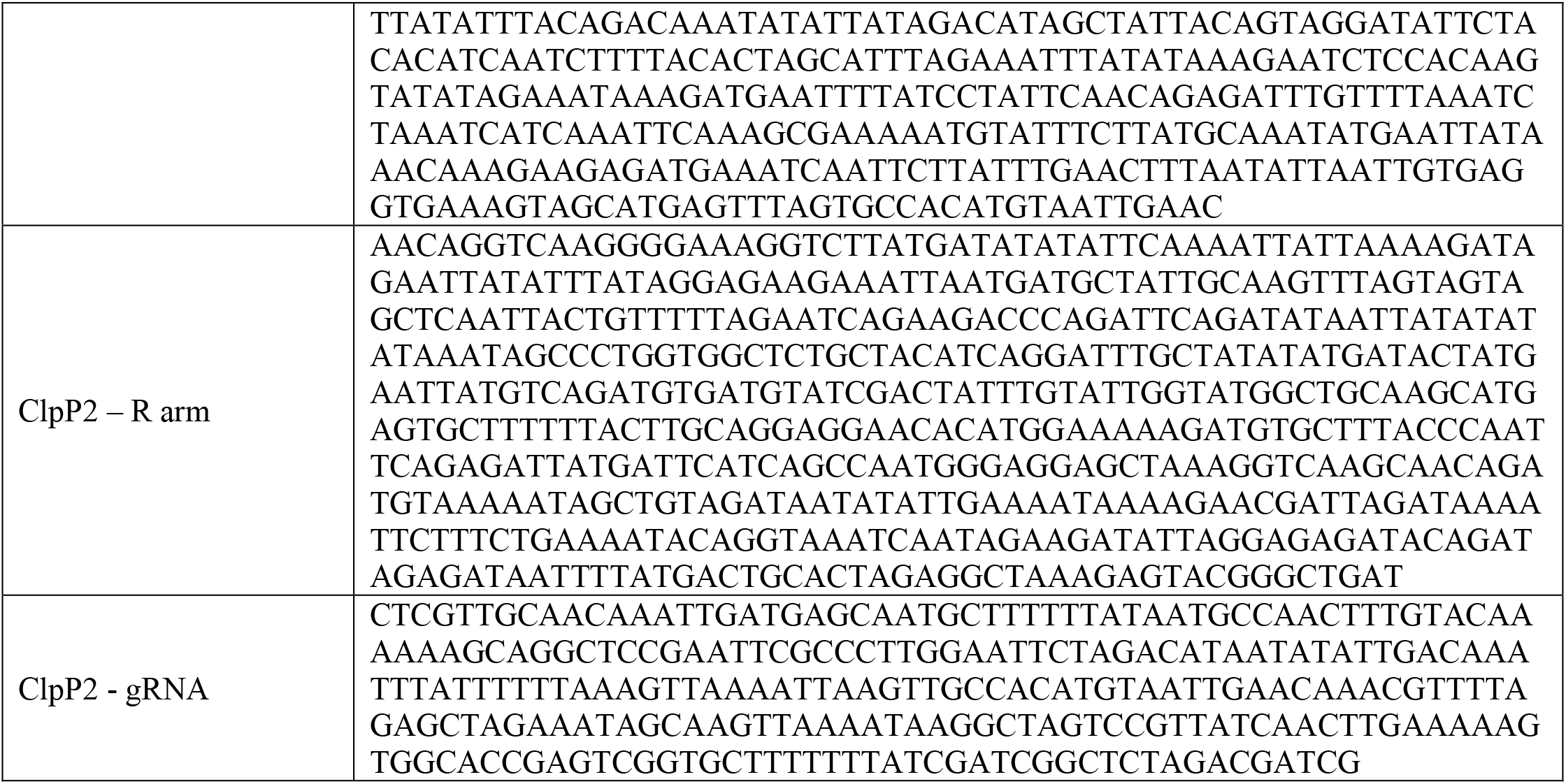
Strains, vectors, oligonucleotides, and CRISPR-Cas9 constructs used in this work.

*E. coli* strains are listed in Table 1 and were grown in Luria-Bertani broth (LB) with shaking at 37°C. Medium was supplemented with chloramphenicol (CAM, 12. 5 μg mL^−1^, ThermoFisher Scientific) and/or kanamycin (KAN, 50 μg mL^−1)^ as indicated.

*B. subtilis* strains are listed in Table 1 and were grown in BHIS broth or on LB agar plates supplemented with CAM (2.5 μg mL^−1)^ and/or tetracycline (TET, 5 and 10 μg mL^−1^, Millipore Sigma).

### Plasmid and strain construction

All primers used in this study are presented in Table 1 and were synthesized by Integrated DNA Technologies (IDT, Coralville, IA). All plasmids and constructed strains are listed in Table 1 as well. The CRISPR-Cas9 nickase vector used for generating *clpP* mutant strains was constructed as previously described (Ahmed, Shadid et al. 2020). For each mutant, three additional parts were designed for insertion into pTMS001: left homology donor template, right homology donor template (**Figure 1A and 1B**), and a single guide RNA (gRNA) (**Figure 1C**). The left and right homology donor templates were designed to insert a NheI restriction site for screening purposes that results in a stop codon early in the *clpP1* and *clpP2* coding regions (**Figure 1D**). The synthetic promoter P4 (Xu, Li et al. 2015) was chosen to drive the expression of customized gRNAs, which produce a single RNA molecule by the fusion of the crRNA and tracrRNA as described by (Doudna and Charpentier 2014). The left and right homology donor templates and the P4::gRNA constructs (Table 1) were synthesized by GenScript and inserted into the NheI and PvuII sites of pTMS001 generating pTMS007 and pTMS008. Sequence verified plasmids (Oklahoma Medical Research Foundation, OMRF) were then transferred into NEB^®^ 10-Beta competent *E. coli* cells via transformation and plated on LB + CAM.

To construct *C. difficile* complementation strains, the vector pTMS005 was constructed as previously described (Ahmed, Shadid et al. 2020) to chromosomally insert *clpP1* or *clpP2*. To generate complement strains, *clpP1* and *clpP2* were PCR amplified from *C. difficile* gDNA using primers Pr-DL0015/16 and Pr-DL0017/18, respectively. pTMSTn was linearized using primers oBAL12F/R and the *clpP1* and *clpP2* fragments were inserted via Gibson assembly, generating pTMS009 and pTMS010, respectively, transferred into NEB^®^ 10-Beta competent *E. coli* cells via transformation, plated on LB + CAM, and sequence-verified. Confirmed vectors were transferred to NEB^®^ Turbo competent *E. coli* cells to generate plasmid multimers and then transferred into *B. subtilis* via transformation (Bouillaut, McBride et al. 2011).

Sequence verified CRISPR-Cas9n and complement plasmids were transferred via transformation into donor strains *E. coli* CA434 or *B. subtilis* BS49 for conjugation with *C. difficile*. For *E. coli* conjugations, 1 mL of an overnight culture of *E. coli* containing the appropriate CRISPR-Cas9n vector was pelleted at 4000 *xg* for 2 min and transferred into the anaerobic chamber. The pellet was resuspended with 200 μL of an overnight *C. difficile* culture. The mixture was spotted on a pre-reduced BHI agar plate (10 x 20 μL spots) and incubated for 24 h at 37°C. The next day the spots were resuspended in 1.5 mL of BHIS and 100 μL was spread onto several BHIS + TAP/D-cycloserine (250 μg mL^−1^, Millipore Sigma) plates. Single colonies were selected and screened via PCR followed by digestion with NheI. Products were run on a 1% agarose gel and screened for the correct digestion pattern followed by sequence verification. Verified *C. difficile* mutants were cured of the vector by serial passaging several times in BHIS alone and PCR screening for the loss of the vector encoded *catP* gene. The *clpP1/clpP2* mutant was constructed as described above by conjugating the pTMS008 vector into the verified *clpP1* mutant.

For *B. subtilis* conjugations, a culture of *C. difficile* was grown overnight in BHIS medium and diluted the next day 1:20 in 5 mL fresh BHIS medium and grown for ~4-6 h. A single colony of *B. subtilis* harboring the appropriate complement vector was inoculated in 5 mL BHIS + CAM/TET and grown aerobically for ~3 h before being passed into the anaerobic chamber. A 1:1 mixture of both cultures was spread onto BHIS plates and incubated at 37°C in the anaerobic chamber. After 24 h, the cells were resuspended in 1.5 mL BHIS and 100 μL was spread onto several BHIS + TAP/KAN plates and incubated for 24 h (TAP selects for integration of the transposon and KAN selects against *B. subtilis*). Single colonies were screened for insertion of *catP* via PCR.

### Growth Curves

Overnight cultures of *C. difficile* strains were diluted to approximately OD_600_ 0.05 in BHIS medium and dispensed into a Bioscreen C plate inside the anaerobic chamber. The plate was sealed, and growth was measured at either 37°C or 42°C using a Bioscreen C plate reader (Growth Curves USA). OD_600_ was measured every 30 min for 24 h.

### Toxicity Studies

Cultures of WT and *clpP* mutants were pelleted at 3,000 xg for 15 min after 72 h growth in order to collect the supernatant. The supernatant was 0.2 μm syringe filtered and serially diluted in the appropriate mammalian cell medium according to cell type (filtering the supernatant removed the bacteria but retained the toxins TcdA and TcdB). All mammalian cell cultures were grown in cell culture-treated plastics in a humidified incubator at 37°C with 5% CO_2_. Caco-2 and CHO cell lines were purchased from ATCC and cultured as recommended by ATCC in Eagles Defined Medium or F-12K medium (Corning), both supplemented with 10% fetal bovine serum. Cell cultures were trypsinized once they had reached ~80% confluence and equal volumes of media were added to neutralize the trypsin. Detached cells were then harvested by centrifugation at 100 xg for 15 min and resuspended in the appropriate medium. 100 μL aliquots (5 x 10^5^ cell density) were used to seed 96-well plates and allowed to grow overnight at 37°C with 5% CO_2_. Once cells had reached 60-70% confluence, the medium was aspirated and serially diluted supernatant from each *C. difficile* strain was added across triplicate wells to the seeded plates. Plates were then incubated with supernatant for 18 h at 37°C with 5% CO_2_ before cell viability was assessed via Cell Counting Kit-8 (CCK-8, Dojindo) according to the manufacturer’s instructions.

### Sporulation Studies

Sporulation and germination of *C. difficile* strains were assessed using methods described previously (Putnam, Nock et al. 2013). All experiments were performed in the anaerobic chamber (6% CO_2_, 3.5% H_2_, and N_2_) unless otherwise noted. Briefly, *C. difficile* strain glycerol stocks were recovered on BHIS plates and incubated overnight at 37°C. Colonies from ~24 h growth on BHIS plates were used to inoculate 2 mL BHIS liquid medium. After ~4 h growth at 37°C, cultures were back-diluted and growth-phase monitored as optical density by measuring absorbance at 600 nm (OD_600_). Once cultures reached an OD_600_ of 0.4 (mid-log phase growth), 120 uL were spread onto 70:30 plates and incubated for ~22 h at 37°C. Cells were harvested into 600 uL of pre-reduced PBS and divided equally into two tubes for the Heat Resistance Assay (HRA). Untreated cells were serially diluted, and dilutions were plated on pre-reduced BHIS-TA plates in triplicate. The heat-treated tubes were exposed to 60°C for 30 min, vortexing every 10 min. Heat-treated cells were then serially diluted and plated on pre-reduced BHIS-TA plates in triplicate. After ~20 h, colony forming units (CFUs) were counted. The percent of heat-treated spores was determined based on the ratio of heat-resistant spores (germinating CFUs from heat-treated samples) to total cells (CFUs from samples that were not heat-treated). Heat-resistance efficiencies were calculated by determining the CFU of germinating heat-resistant spores for mutant strains compared to WT. Results are based on a minimum of three biological replicates.

Within 1 h of harvesting, aliquots of the un-treated samples were used for phase-contrast microscopy spore counts as described previously (Burns and Minton 2011, Putnam, Nock et al. 2013). Briefly, samples were washed once with PBS and the pellet was resuspended in 190 uL of PBS. Sporulation was assessed by embedding 5 uL of sample into 1% agarose gel pads, sealed with a coverslip, and assessed in replicates of 4 or more per strain on an Olympus CX41RF compound microscope (Olympus America Inc.) equipped with phase objectives. Images were captured with the Olympus SC50 5 mp digital USB CCD detector and processed with CellSens software (Olympus America Inc.). Spores and vegetative cells were counted using ImageJ software (Schneider, Rasband et al. 2012).

### Transmission Electron Microscopy

The OMRF Imaging Core Facility performed TEM on samples as described herein. Spores were fixed with 4% Paraformaldehyde (EM grade) and 2.5% Gluteraldehyde (EM grade) in 0.2 M sodium cacodylate buffer for 1 day at 4°C. Samples were then post fixed for 60 min in 1% osmium tetroxide (OsO_4_) in 0.2 M sodium cacodylate and rinsed 3X for five minutes each in 0.2 M sodium cacodylate buffer. The samples were stored in 0.2 M sodium cacodylate buffer overnight on a rocker at room temperature. The samples were then dehydrated in a graded ethanol series (50%, 60%, 75%, 85%, 95%, and 100%) with rocking incubation for 15 min at each ethanol concentration. Then the samples had two 15-min treatments in 100% propylene oxide. Following dehydration, the samples were infiltrated in a graded epon/araldite (EMS) resin/propylene oxide series at ratios of 1:3, 1:1, and 3:1 for 60 min, overnight, and 120 min the next day, respectfully. The spores were further infiltrated with pure resin for 45 min, 90 min, and then overnight. The samples were embedded in resin plus BDMA (accelerator) and polymerized at 60°C for 48 h. Ultrathin sections were stained with Sato’s lead and saturated uranyl acetate in 50% methanol before viewing on a Hitachi H7600 Transmission Electron Microscope.

All spore counts and measurements were performed using ImageJ software (Schneider, Rasband et al. 2012). Spores with a gray/dark spore core or a bright spore core and atypical spores were counted for triplicate TEM grid images. A total of at least 50 structures were counted for each image except for the *clpP1* mutant as a total of 50 spores were not present on any one image; however, across a total of three images >50 spores were counted (n = 103) for the *clpP1* mutant as well (Supplemental Table S1). A spores was considered “typical” if it had at least a well-defined spore core, cortex, and coat. Any structures that did not have an electron dense spore coat nor other well-defined features (e.g. missing apparent cortex) were considered “atypical” spores. The total percent spores with gray/dark cores was calculated by dividing the total number of spores with gray/dark cores by the total number of typical spore structures and multiplying by 100. The same was done for spores with bright cores. The total percent atypical spores was calculated by dividing the total number of atypical spores by the total number of structures counted and multiplying by 100. The same was done for typical spores. The spore core, cortex, and coat were measured in ImageJ as an average of four different measurements on each side of the spore for 50 total typical spores. These averages were compiled in Supplemental Table S2.

### Sample Preparation and Proteomic Analysis

*difficile* strain 630 and the *clpP* mutants thereof were grown for 24 h on BHIS solid medium in an anerobic chamber at 37°C. Colonies (~10) were transferred to liquid BHIS, incubated for 5 hours at 37°C, back diluted 1:50 in liquid BHIS and allowed to reach exponential growth by incubating at 37°C until samples reach an OD (600 nm) of 0.4. Cultures (120 uL) were plated on BHIS and incubated at 37°C for 48 hours before harvesting. The entire plate was harvested into 1000 uL PBS before removing samples from the anaerobic chamber. The cells were washed 1X with PBS. Samples were lysed with freshly prepared 8 M urea lysis buffer and centrifuged. Protein concentration was measured (Pierce BCA Protein Assay, Thermo Fisher Scientific, IL, USA) and a total of 100 μg of protein per sample was subjected for overnight trypsin digestion. Tryptic peptides were desalted using C18 Sep-Pak plus cartridges (Waters, Milford, MA) and were lyophilized for 24 hours to dryness. The dried peptides were reconstituted in buffer A (0.1 % formic acid) at a concentration of 0.25 μg/μl and an equal amount of proteins (1 μg) was injected for LC-MS/MS analysis.

The LC-MS/MS was performed on a fully automated proteomic technology platform that includes a Dionex UltiMate ® 3000 (Thermo Fisher Scientific, USA) system connected to a Q Exactive HF-X mass spectrometer (Thermo Fisher Scientific, Waltham, MA). The LC-MS/MS set up was slightly modified from the original set up described earlier (Kelstrup, Bekker-Jensen et al. 2018, Bian, Zheng et al. 2020). Peptides were loaded onto an in-house packed trap column (150 μm × 3 cm, packed with Bio-C18 3 μm resin, Sepax Technologies, DE, USA) with solvent A (0.1% formic acid in LC-MS grade water) at a flow rate of 3 μl/min for 10 min and separated on an analytical column (75 μm × 30 cm, packed in house with Bio-C18 3 μm resin, Sepax Technologies) at 350 nl/min. The analytical column was heated to 55 °C and the peptides were separated through a 70 min-long linear gradient from 0% to 40% buffer B (0.1 % formic acid in acetonitrile). Nanoelectrospray was obtained using a fused silica emitter (PicoTip emitter, New Objective, Woburn, MA). The total run time was of 90 min including column wash and re-equilibration.

The MS/MS analysis was optimized from the earlier published protocol (Kelstrup, Bekker-Jensen et al. 2018). Briefly, the Q Exactive HF-X mass spectrometer was operated as follows: Positive polarity; spray voltage 2.0 kV, funnel RF lens value at 40, and capillary temperature of 320 °C. The data dependent acquisition (DDA) using the Full MS-ddMS2 setup was used. The spectra were collected using a top-15 data-dependent method. Full MS resolution was set to 60,000 at m/z 200 and full MS AGC target was 3E6 with a maximum injection time (IT) of 45 ms. Mass range was set to 350–2000. AGC target value for fragment spectra was set to 1E5. The isolation width was set to 1.3 m/z, and the first mass was fixed at 100 m/z. The normalized collision energy was set to 28%. Peptide match was set to preferred, and isotope exclusion was on. MS1 and MS2 spectra were acquired in profile and centroid mode, respectively. Precursors were fragmented by HCD using 30% normalized collision energy (NCE) and analyzed in the Orbitrap at a resolution of 15,000. The dynamic exclusion duration of fragmented precursor ions was set to 30 s. The ion selection abundance threshold was set at 2E5 with charge state exclusion of unassigned and z =1, or 6-8 ions. Peptide spectrum matching of MS/MS spectra of each file was searched against the NCBI *C. difficile* 630 (TaxonID: 272563, downloaded on 10/06/2020) and *C. difficile* R20291 (TaxonID: 645463, downloaded on 10/06/2020) databases using the Sequest algorithm within Proteome Discoverer v 2.4 software (Thermo Fisher Scientific, San Jose, CA). The Sequest database search was performed with the following parameters: trypsin enzyme cleavage specificity, 2 possible missed cleavages, 10 ppm mass tolerance for precursor ions, 0.02 Da mass tolerance for fragment ions. Search parameters permitted dynamic modification of methionine oxidation (+15.9949 Da) and static modification of carbamidomethylation (+57.0215 Da) on cysteine. Peptide assignments from the database search were filtered down to a 1% FDR. The relative label-free quantitative and comparative among the samples were performed using the Minora algorithm and the adjoining bioinformatics tools of the Proteome Discoverer 2.4 software. All identification and quantification protein data are available in Supplementary Table S3.

## Author Contributions

C.E.B., T.S., N.P.L., and M.L.K. conducted the experiments and acquired the data pertaining to mutant generation and behavior. N.A. conducted the proteomic analysis and authored the methods and materials and sections of the manuscript pertaining to this work. C.E.B. N.P.L. and A.S.D. designed the research studies and decided the trajectory of the research. J.D.B. provided critical insight, expertise, personnel, and facilities, and reviewed the manuscript. C.E.B. and N.P.L. wrote the narrative. All authors reviewed and accepted the final version of the manuscript.

## Acknowledgements

The authors gratefully acknowledge Drs. Aimee Shen, Shonna McBride, and Joseph Sorg for providing insightful discussion during our TEM initial studies. Additionally, we thank Dr. McBride for sharing with us the vectors needed for complementation and Dr. Sorg for sharing the *Bacillus* conjugal strain used in these studies. Research reported in this publication was supported by an Institutional Development Award (IDeA) from the National Institute of General Medical Sciences of the National Institutes of Health under grant number P20GM103640 (A.S.D.) and National Institute of Allergy and Infectious Diseases of National Institutes of Health grant number R01AI119048 (J.D.B.). N.A. gratefully acknowledges the OU VPRP Office for the initial funding support for the establishment of the Proteomics Core Facility. The content is solely the responsibility of the authors and does not necessarily represent the official views of the National Institutes of Health.

